# Thermal boundaries and underlying systems biology mechanisms in two waterbloom cyanobacteria

**DOI:** 10.1101/2025.11.24.690146

**Authors:** Tainyue Zhang, Li Liu, Zeshuang Wang, Ziyan Zhao, Yiyue Guo, Huansheng Cao, Zhou Yang

## Abstract

Climate change and recent heatwaves have elevated temperature to the level of nutrients in driving CHABs. In this new dimension of CHABs ecology, it is important to understand the thermal boundaries of each blooming species. One well-known example is the temporal and thermal partitioning between filamentous, nitrogen-fixing taxa such as *Dolichospermum* spp. and *Pseudanabaena* spp., which dominate in spring, and colonial/coccoid *Microcystis aeruginosa*, which predominates in summer and autumn. Here we show that *Dolichospermum* sp. and *M. aeruginosa* display converse temperature-growth relations, *Dolichospermum* sp. grows better at low temperatures (13-15°C) and *M. aeruginosa* grows better at high temperatures (29-31°C), with similar growth rates between 18 and 25°C. This thermal separation is sustained by systems-level changes, in terms of the macromolecule contents—proteins, ATP, carbohydrates, and lipids—and transcriptome and metabolic fluxes to critical unsaturated fatty acids, which were experimentally verified by supplementing these FAs. The gene regulatory and systems biology mechanisms are explored. Through homologous and evolutionary analyses, a conserved Hik34–Rre1–RpoD cascade underlying differential temperature-responsive gene regulation. Enzyme-constrained metabolic models reproduced genus-specific lipid shifts and uncovered contrasting flux reorganizations under thermal stress. Together, these results establish the metabolic and regulatory foundations that shape the distinct thermal niches of bloom-forming cyanobacteria, providing mechanistic insight into how climate warming may restructure their seasonal dominance.

## Introduction

Cyanobacterial harmful algal blooms (CHABs) represent one of the most pervasive ecological hazards of the Anthropocene, notable for their global expansion, increasing intensity, and the persistent difficulty of mitigation. First recognized in the 1870s for their toxicity to animals ^1,2^, CHABs have since been examined for more than a century, leading to broad consensus that eutrophication—whether chronic or episodic ^3^ — is the primary driver of bloom formation. More recently, the functional traits underpinning cyanobacterial dominance have been systematically delineated ^4^ and integrated into the CyanoPATH database ^5^. These advances have led to a proposed two-tier mechanism in which interactions between ecophysiology and environmental conditions operate across population and community scales ^6^.

Beyond nutrient enrichment, temperature has long been recognized as a key driver of CHAB dynamics ^7^. Contemporary climate change—particularly the increasing frequency and severity of heatwaves ^8–10^ —has elevated thermal forcing to a second major determinant of bloom formation, comparable in influence to continued nutrient inputs. Under this emerging thermal dimension of CHAB ecology, delineating the temperature boundaries of individual bloom-forming species has become essential for improving bloom prediction and guiding effective management strategies. A well-documented example is the temporal and thermal partitioning between filamentous, nitrogen-fixing taxa such as *Dolichospermum* spp. and *Pseudanabaena* spp., which dominate in spring, and colonial/coccoid *Microcystis aeruginosa*, which predominates in summer and autumn ^9,11,12^. From a systems-biology perspective, these thermal boundaries reflect the integrated responses of genome-scale metabolic networks (GSMs) to environmental temperature ^13–15^.

In this study, we sought to define the thermal ranges and growth optima of *Dolichospermum* spp. and *M. aeruginosa*, and to elucidate the systems-level mechanisms that govern their thermal boundaries and temperature-dependent growth.

## Results

### Optimal temperatures for growth and intracellular macromolecule contents

Three major differences in thermal responses were evident between *Dolichospermum* sp. and *Microcystis aeruginosa*. First, the two species exhibited contrasting temperature–growth relationships (Fig. 1a). At lower temperatures (13–15 °C), *Dolichospermum* sp. strains grew substantially faster than *M. aeruginosa*. Between 15 and 25 °C, growth rates of the two species converged, with *Dolichospermum* sp. reaching its maximal growth at 25 °C. In contrast, *M. aeruginosa* achieved its highest growth rates at 29–32 °C, markedly exceeding those of *Dolichospermum* sp. at these temperatures.

**Figure 1.**
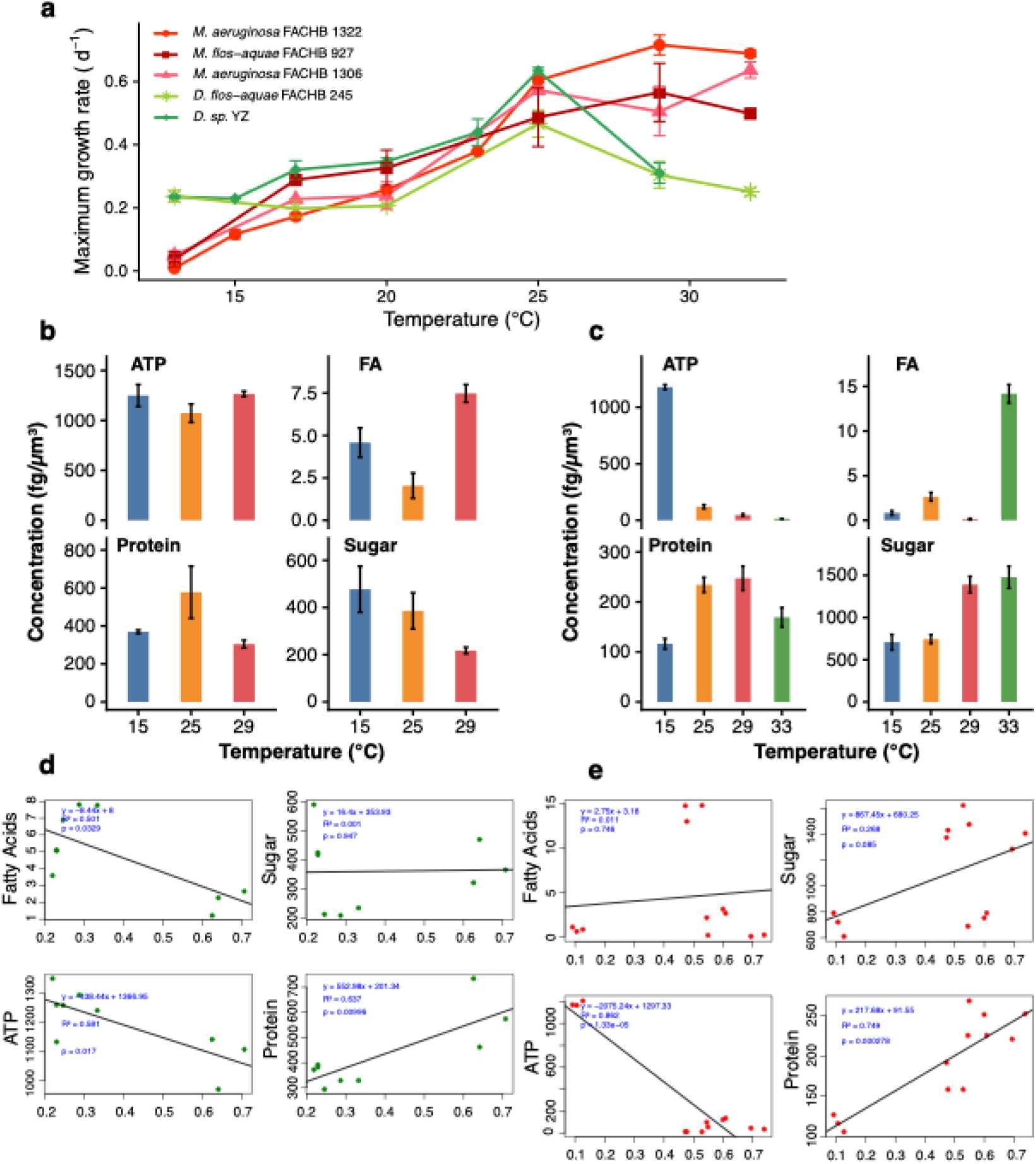
Temperature-dependent growth and macromolecular composition. **(a)** Maximum growth rates of *Microcystis aeruginosa* (M) and *Dolichospermum* sp. (D) across temperatures. **(b)** Cellular ATP, carbohydrates, total protein and fatty acid contents of *Dolichospermum* sp. at 15, 25, 29°C. **(c)** Cellular ATP, carbohydrates, total protein and fatty acid contents of *M. aeruginosa* at 15, 25, 29 and 33 °C. **(d)** Regression relationships between growth rate and ATP, sugar, protein and fatty acid contents in *Dolichospermum* sp. Regression relationships between growth rate and ATP, sugar and protein contents in *M. aeruginosa*.

Next, the contents of four major macromolecules—ATP, lipids, proteins, and carbohydrates—were quantified at selected temperatures. Intracellular contents were temperature-dependent and differed between the two species. In *Dolichospermum* sp. at 15, 25, and 29 °C, ATP and fatty acids reached their lowest levels at 25 °C, whereas protein content peaked at this temperature, and carbohydrate content declined persistently. In *M. aeruginosa* at 15, 25, 29, and 32 °C, protein content showed a similar trend to *Dolichospermum* sp., increasing with temperature and declining at the highest temperature. Fatty acids displayed a variable pattern, rising from 15 to 25 °C, declining at 29 °C, and increasing sharply at 32 °C. ATP and carbohydrates exhibited opposite responses: ATP levels decreased steadily with increasing temperature, whereas carbohydrate content rose consistently.

To further assess how macromolecular patterns relate to growth, regression analysis was performed between growth rate and the contents of major macromolecules. The analysis revealed consistent correlations of ATP and protein levels with growth in both species: higher growth rates were associated with increased protein but decreased ATP. Sugar content showed a positive correlation with growth in *M. aeruginosa*, whereas in *Dolichospermum* sp. no significant relationship was observed. Fatty acids showed opposite trends between the two species: in *Dolichospermum* sp., faster growth corresponded to lower fatty-acid abundance, whereas in *M. aeruginosa*, higher growth rates were associated with slightly elevated fatty-acid levels (Fig. 1d,e).

### Lipid Composition and ^13^C-Based Carbon Flux to Fatty Acids

Given differences in intracellular lipid contents with respect to temperature and growth rate, fatty-acid composition and ^13^C-based enrichment were examined to explain species-specific temperature optima. *M. aeruginosa* contained three long-chain polyunsaturated fatty acids (PUFAs)—C18:3n6(6,9,12), C20:3n6(8,11,14), and C22:1T—not detected in *Dolichospermum* sp., whereas *Dolichospermum* sp. contained one monounsaturated fatty acid (MFA), C14:1, absent in *M. aeruginosa*.

All fatty acid contents were generally lowest at peak growth rates in both species (Fig. 2a). Fatty acids most abundant at low temperatures were largely similar between species, with a few exceptions (C18:3n6(9,12,15) and C14:0), which reached their highest levels in *Dolichospermum* sp. at 29 °C. At high temperature, both MFAs and PUFAs were abundant in *M. aeruginosa*, some of these fatty acids were also present in *Dolichospermum* sp.at 29 °C (C18:1n9c(9), C18:2n6c(9,12), C16:0, and C16:1(9)).

**Figure 2.**
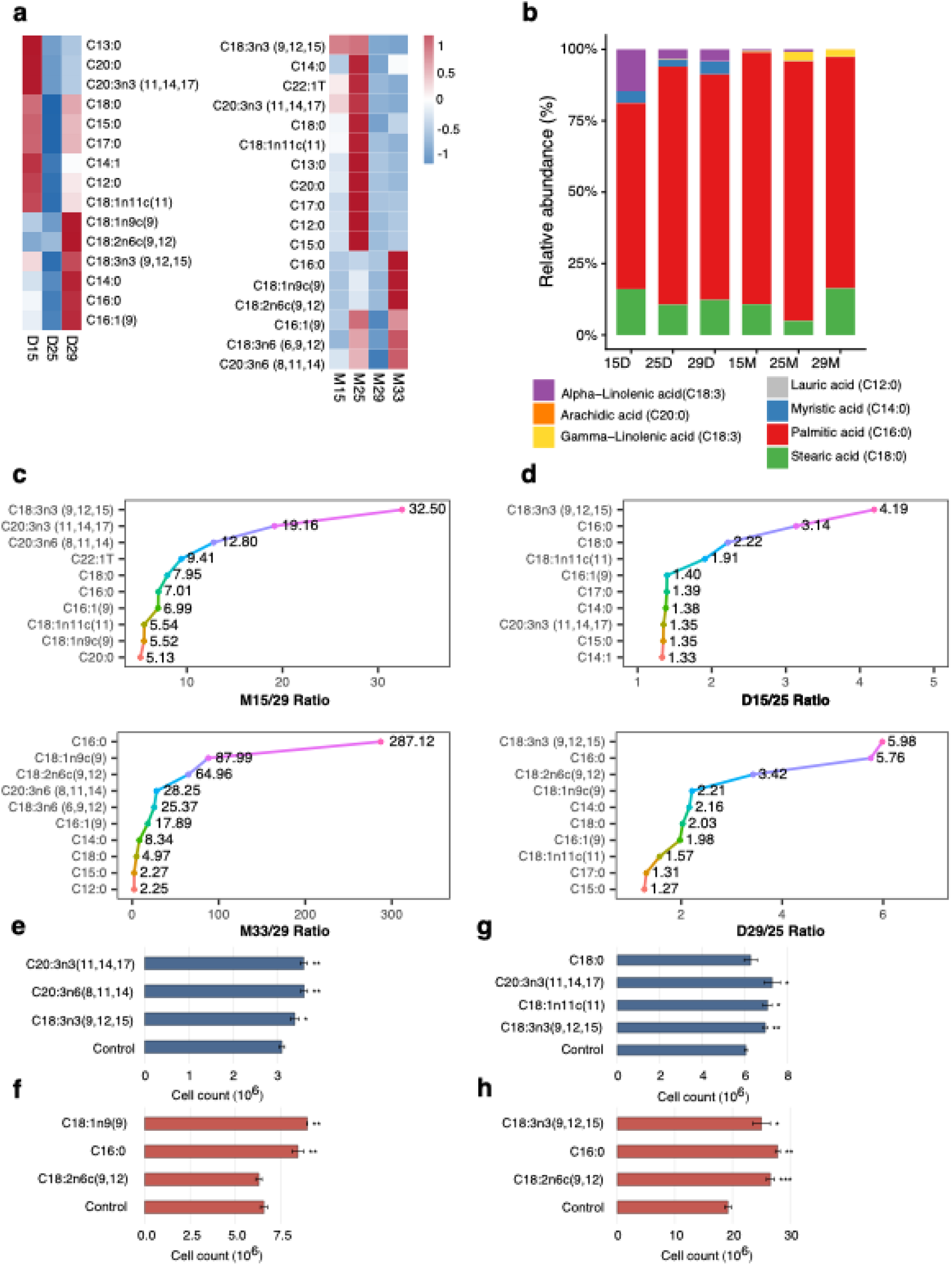
Temperature-dependent ^13^C incorporation, fatty acid enrichment, and functional verification. **(a)** Heatmap of fatty acid abundance in *Dolichospermum* sp. and *M. aeruginosa* across temperatures. **(b)** Relative ^13^C enrichment under different temperatures. **(c)** Top 10 fatty acids in *M. aeruginosa* showing the highest fold changes at 15 °C or 33 °C relative to the optimal temperature (29 °C). **(d)** Top 10 fatty acids in *Dolichospermum* sp. showing the highest fold changes at 15 °C or 29 °C relative to the optimal temperature (25 °C). **(e,f)** Cell numbers of *M. aeruginosa* after 5 days of growth under low temperature (e) or high temperature (f), with or without supplementation of the corresponding exogenous fatty acids. **(g,h)** Cell numbers of *Dolichospermum* sp. after 5 days of growth under low temperature (g) or high temperature (h), with or without supplementation of the corresponding exogenous fatty acids.

To further link these compositional shifts to carbon allocation, ^13^C-labeling experiments were conducted. They revealed strong temperature-dependent redistribution of carbon among fatty acids (Fig. 2b).At their respective optimal temperatures, ^13^C was primarily incorporated into C16:0, followed by C18:0. In *Dolichospermum* sp., C18:3 (α-linolenic acid) and C14:0 were relatively more enriched, whereas these two fatty acids were scarce in *M. aeruginosa*.

Conversely, *M. aeruginosa* showed higher ^13^C enrichment in C18:3 (γ-linolenic acid) compared with *Dolichospermum* sp. In *Dolichospermum* sp., C18:0, C14:0, and C18:3 (α-linolenic acid) were enriched at both low and high temperatures relative to 25 °C. In *M. aeruginosa*, low temperatures increased the fractional ^13^C abundance of C16:0, while C18:0 declined.

Next, the ratios of fatty-acid abundance at non-optimal temperatures relative to the optima were calculated to identify those that support growth under thermal stress (Fig. 2c,d).Fatty acids with high ratios at low temperatures but low ratios at high temperatures were inferred to facilitate cold growth, such as C18:3n3(9,12,15) and C20:3n3(11,14,17) in *M. aeruginosa*. Fatty acids with high ratios at high temperatures but low ratios at low temperatures were considered to support growth under heat stress, including C16:0, C18:1n9(9), and C12:2n6(9,12) in *M. aeruginosa*, and C12:2n6(9,12) in *Dolichospermum* sp. Some fatty acids in *Dolichospermum* sp., such as C18:3n3(9,12,15) and C16:0, exhibited high ratios at both low and high temperatures, suggesting a broader role in supporting growth under thermal stress.

To verify these predictions, the top 3–4 fatty acids at each non-optimal temperature were added exogenously to cultures for five days at concentrations corresponding to measured intracellular abundances. Cell densities were monitored daily. In *M. aeruginosa* (Fig. 2e,f), only C18:2n6c(9,12) maintained control-like growth, whereas all other supplemented fatty acids significantly increased cell numbers. In *Dolichospermum* sp. (Fig. 2g,h), only low-temperature C18:0 did not differ significantly from controls, although values were slightly elevated; all other additions enhanced growth. Consistently, Supplementary Fig. 1a,b show that under low and high temperatures, most supplemented groups achieved higher maximum growth rates than controls. These results indicate that specific fatty acids can partially rescue or enhance growth under thermal stress, with species- and temperature-specific differences in effectiveness.

### Transcriptomic Responses to Temperature

Gene expression at low and high temperatures was compared to relative optimal temperature, 29°C in *M. aeruginosa* and 25°C in *Dolichospermum* sp., and significantly up- or downregulated genes were identified for functional enrichment analysis. GO-term enrichment revealed temperature-dependent functional shifts (Supplementary Fig. 1c–j).

At low temperature, *M. aeruginosa* upregulated general metabolic and protein-biosynthesis pathways, while downregulating nucleotide biosynthesis, DNA replication, NADPH generation, and acetyl-CoA production. In contrast, *Dolichospermum* sp. upregulated photosynthesis, electron transport, and cytokinesis-related pathways at 15 °C, while downregulating NADPH generation, acetyl-CoA biosynthesis, protein folding, and specific nitrogen-metabolism pathways.

At high temperature, *M. aeruginosa* upregulated pathways related to cell shape, cell-wall biogenesis, and amino-acid biosynthesis—including glutamine and arginine—while downregulating photosynthesis. Conversely, *Dolichospermum* sp. upregulated photosynthesis, protein folding, and energy-related processes (carbon fixation, oxidative phosphorylation, and ATP-linked electron transport) at 29 °C, while downregulating general metabolism, gene expression, and peptide biosynthesis.

Based on GO-term enrichment results, genes from major pathways involved in central carbon metabolism, energy supply, and fatty-acid biosynthesis were visualized as heatmaps (TPM; Fig. 3) to examine their temperature-dependent expression patterns. Gene expression in oxidative phosphorylation, fatty-acid biosynthesis, glycolysis, and the pentose-phosphate pathway differed markedly between species, whereas genes in the calvin cycle and photosynthesis showed similar expression patterns, peaking at their respective optimal temperatures. In *Dolichospermum* sp., fatty-acid biosynthesis genes peaked at 15 °C, whereas in *M. aeruginosa*, they peaked at 33 °C. Genes involved in glycolysis, the TCA cycle, the pentose-phosphate pathway, and oxidative phosphorylation generally peaked at low temperature in *Dolichospermum* sp., but at high temperature in *M. aeruginosa*.

**Figure 3.**
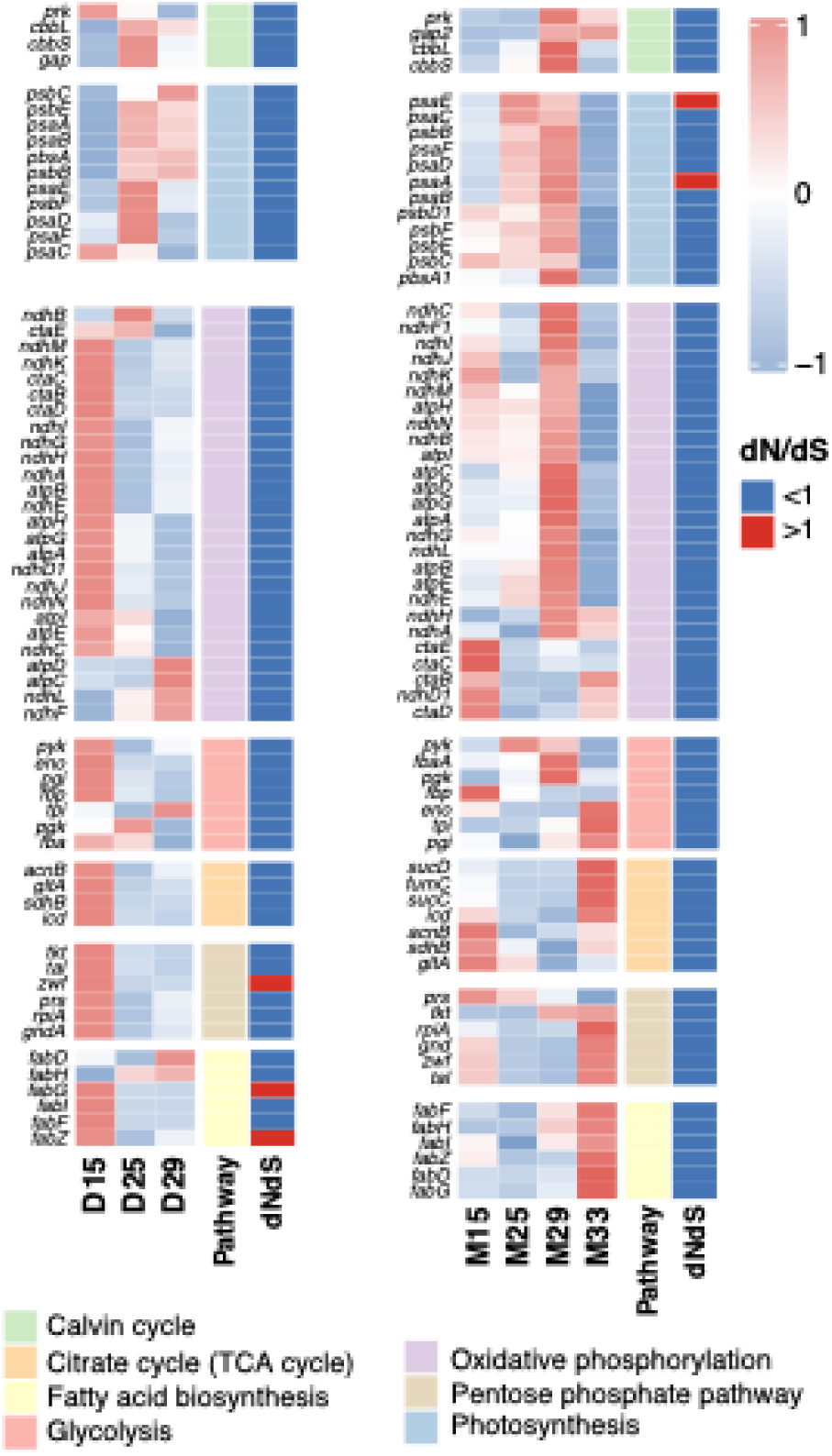
Temperature-dependent transcriptional responses and evolutionary constraints. Heatmap of gene expression (TPM) across temperatures for major metabolic pathways in *Dolichospermum* and *Microcystis aeruginosa*, with accompanying dN/dS ranges. Genes with dN/dS > 1 are highlighted in red.

To explore the evolutionary conservation of these temperature-responsive genes, dN/dS were analyzed and revealed lineage-specific selective pressures on these temperature-responsive pathways (Fig. 4a,b). Several genes in fatty-acid biosynthesis and the pentose-phosphate pathway showed dN/dS > 1 in *Dolichospermum* sp., whereas two photosynthesis-related genes exceeded 1 in *M. aeruginosa*, suggesting divergent selection shaping the evolution of metabolic responses to temperature.

**Figure 4.**
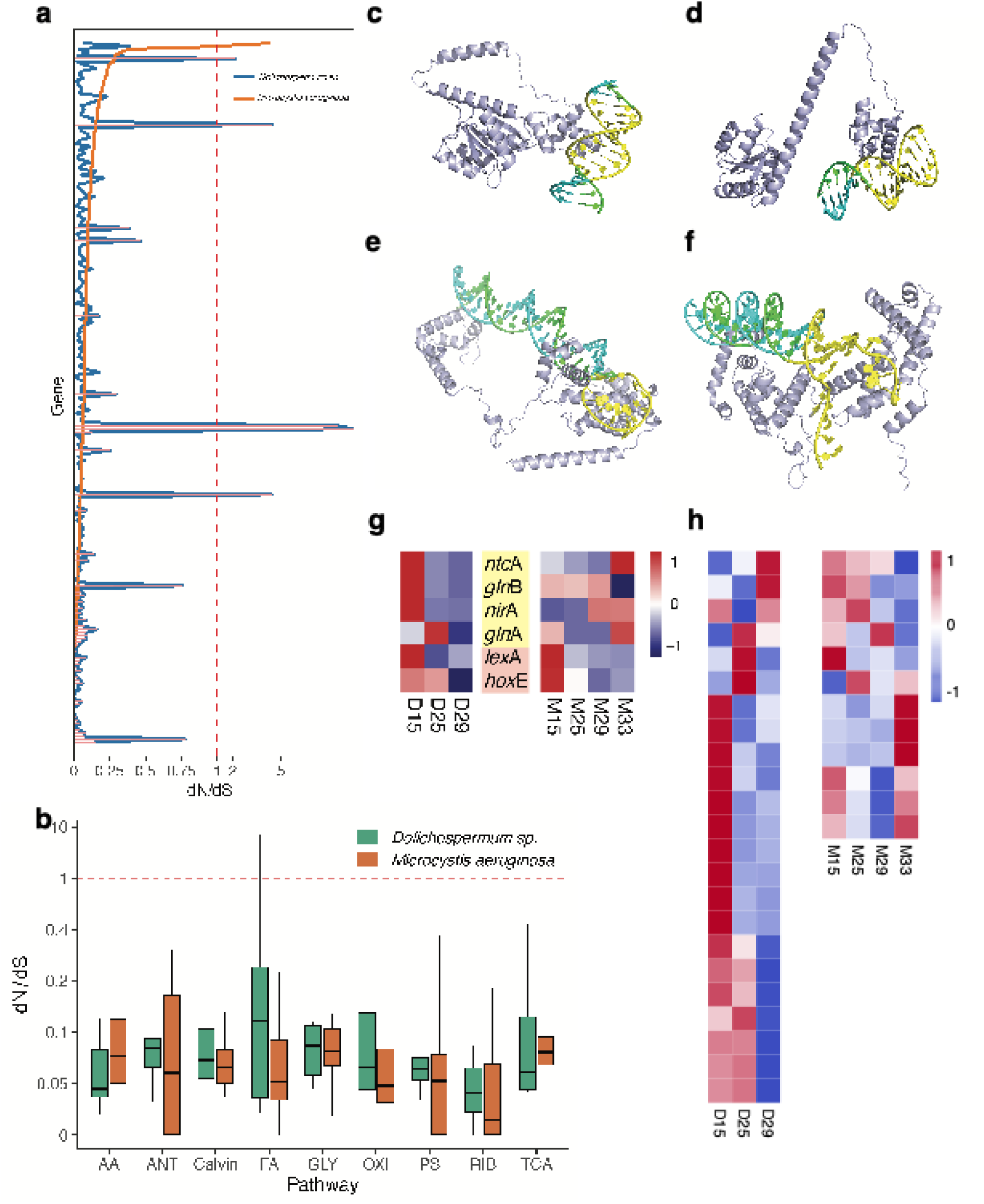
Evolutionary divergence, transcriptional regulation and structural predictions. (a) Differences in dN/dS values of genes in major metabolic pathways between *Dolichospermum* sp.and *M. aeruginosa* . (b) Box plots comparing dN/dS values of genes across major metabolic pathways between *Dolichospermum* sp. and *M. aeruginosa*. Pathway abbreviations are as follows: AA, biosynthesis of amino acids; Calvin, Calvin cycle; TCA, citrate cycle (TCA cycle); OXI, oxidative phosphorylation; FA, fatty acid biosynthesis; GLY, glycolysis; PS, photosynthesis; ANT, photosynthesis–antenna proteins; RIB, ribosome. (c) AlphaFold3-predicted complex between the transcription factor Rre1 (WP_028090035.1) and the promoter region of *rpoD* (ACIR131C_RS0104995) in *Dolichospermum* sp.. (d) AlphaFold3-predicted complex between Rre1 (ELP54679.1) and the promoter region of *rpoD* (O53_4892) in *M. aeruginosa*. (e) AlphaFold3-predicted complex between RpoD (WP_028089309.1, ACIR131C_RS0104995) and the promoter region of ACIR131C_RS0101185 in *Dolichospermum* sp.. (f) AlphaFold3-predicted complex between RpoD (ELP53163.1, O53_4892) and the promoter region of O53_4547 in *M. aeruginosa*. (g) Heatmaps of expression for known transcription factors NtcA and LexA and their associated target genes (yellow background for NtcA regulon genes: *glnB*, *nirA*, *glnA*; red background for LexA regulon gene: *hoxE*) in *Dolichospermum* sp. and *M. aeruginosa* across temperatures. Heatmaps showing expression of genes predicted by AlphaFold3 to be potential RpoD targets in *Dolichospermum* sp. (left) and *M. aeruginosa* (right) across temperatures.

### Systems-level gene regulation and operation in metabolic network sustaining optimal growth in two species

The distinct thermal preferences and associated metabolite support underpin niche separation between *Dolichospermum* sp. and *M. aeruginosa* observed in nature. Mechanistically, these differences arise from systems-level metabolic regulation, reflected in divergent transcriptional programs across temperature regimes. To identify the regulatory hierarchy shaping these responses, we examined both established and putative transcriptional regulators in each species.

Key TFs exhibited contrasting temperature-dependent expression patterns. In *Dolichospermum* sp., the nitrogen-regulatory factor NtcA and its targets (*glnB* and *nirA*), as well as the global stress-response regulator LexA with *hoxE*, showed maximal expression at 15 °C. In *M. aeruginosa*, by contrast, NtcA and its associated genes (*glnB* and *nirA*) peaked at higher temperatures (29–33 °C), whereas *lexA* and *hoxE* peaked at low temperature, as observed in Dolichospermum sp. (Fig. 4g).

Here we focus on a putative thermal sensor histidine kinase Hik34, which upon activation phosphorylates effector protein Rre1 in *Synechococcus elongatus* PCC 7942^16^. One homolog was identified in *Dolichospermum* sp. and *M. aeruginosa*, with sequence identity of WP_028090232.1 and ELP53887.1. To further confirm these homologs are conserved functional sensor kinases, we first verified their evolutionary conserveness, i.e., under negative selection (minimizing sequence divergence). Molecular evolution analyses showed that these two genes are among the most conserved, with dN/dS ratios of hik34 = 0.178, 0.1776, and rre1 = 0.0744, 0.2245 across *Dolichospermum* sp., and *M. aeruginosa* (Fig. 4a).These genes were conserved at the same levels as those genes involved in core metabolism (Fig. 4b).

Next, we tested if these two homologs were kinases by expressing both sensor and effector proteins in E coli and quantifying ADP levels from phosphorylation of Rre1 by Hik34 in vitro.

Rre1 in *E. coli* has been shown to bind the promoter regions sigma factor *σ*^70^ (encoded by *rpoD*)^17^. Here using AlphaFold3^18^, we confirm this is also the case: Rre1 showed high-affinity binding to the *rpoD* promoter regions in *Dolichospermum* sp. and *M. aeruginosa* (Fig. 4c, d) with ipTM and pTM being 0.73, 0.52 (in *Dolichospermum* sp.), and 0.63,0.53 (in *M. aeruginosa*). Furthermore, *σ*^70^ as one of the major sigma factors of the RNA polymerase holoenzyme, is shown to bind was predicted to bind the promoter regions of multiple downstream genes at two very conserved hexamers -35 TTGACA and -10 TATAAT in *E. coli*^17^. AlphaFold3 predicted ACIR131C_RS0101185 and O53_4547 genes bound to their respective *σ*^70^ in *Dolichospermum* sp. and *M. aeruginosa*, with confidence (ipTM > 0.8 and pTM >0.5). Fig. 4e,f illustrating one representative high-confidence example in each species. These models suggest that phosphorylated Rre1 by sensor kinase Hik34 acts upstream of *rpoD* to regulate its transcription, and expressed *σ*^70^ in turn directly controls the expression of downstream stress-responsive target genes.

Genes with predicted RpoD binding displayed distinct genus-level temperature responses (Fig. 4h). In *Dolichospermum*, most RpoD-associated genes peaked at low temperature, with fewer at optimal or high temperatures; in *M. aeruginosa*, roughly half peaked at high temperature, while a smaller subset peaked at low temperature.

### Metabolic Flux Analysis of Fatty-Acid Biosynthesis

Finally, we integrated transcriptome, lipidome, gene regulation, and enzymatic parameters into a genome-scale metabolic network to simulate the systems biology response in each species. To that end, we first reconstructed the GSMs, *i*Dfs703 for *Dolichospermum* sp. and *i*Mar695 for *M. aeruginosa* from their genomes, respectively (Table 1). *i*Dfs703 has 703 genes encoding enzymes and 1,280 reaction, and 1,286 metabolites; *i*Mar695 has 695 genes encoding enzymes and 1,202 reaction, and 1,225 metabolites. Biomass compositions were also provided by experimentally measuring the total protein content, sugar, ATP, and fatty acids. Each GSM was further converted to Gecko model, which treats enzymes as metabolites to facilitate adjusting enzymes-related parameters^19^.

**Table 1.** Parameters for the genome-scale metabolic networks (GSMs) of *Dolichospermum* sp. and *Microcystis aeruginosa*.

|  | *iDfs703* | *iMar695* |
| --- | --- | --- |
| *Included genes* | 703 (14%) | 695(14%) |
| *Reactions* | 1,280 | 1,202 |
| *Metabolic reactions* | 1,117 | 1,068 |
| *Transport reactions* | 85 | 70 |
| *Exchange reactions* | 78 | 64 |
| *Metabolites* | 1,286 | 1,225 |
| Unique metabolites | 1,213 | 1,164 |
| Cytoplasmic | 1,286 | 1,225 |
| Extracellular | 73 | 61 |
| *Gene-protein-reaction associations* |  |  |
| Gene associated (metabolic/transport) | 1,032 | 960 |
| No gene association (metabolic/transport) | 248 | 172 |

By creating condition-specific GSMs by incorporating transcriptomes and adjusting enzyme parameters at each temperature, metabolic-flux modeling revealed divergent temperature-dependent fatty-acid biosynthetic strategies consistent with lipidomic patterns. Fluxes of fatty-acid biosynthetic reactions at optimal temperatures showed substantial differences between *Dolichospermum* sp. and *M. aeruginosa* (Fig. 5a). Specifically, the fluxes were higher in *Dolichospermum* sp. than *M. aeruginosa* at their respective optima, consistent with temperature-dependent fatty-acid abundance patterns (Fig. 1b, c).

**Figure 5.**
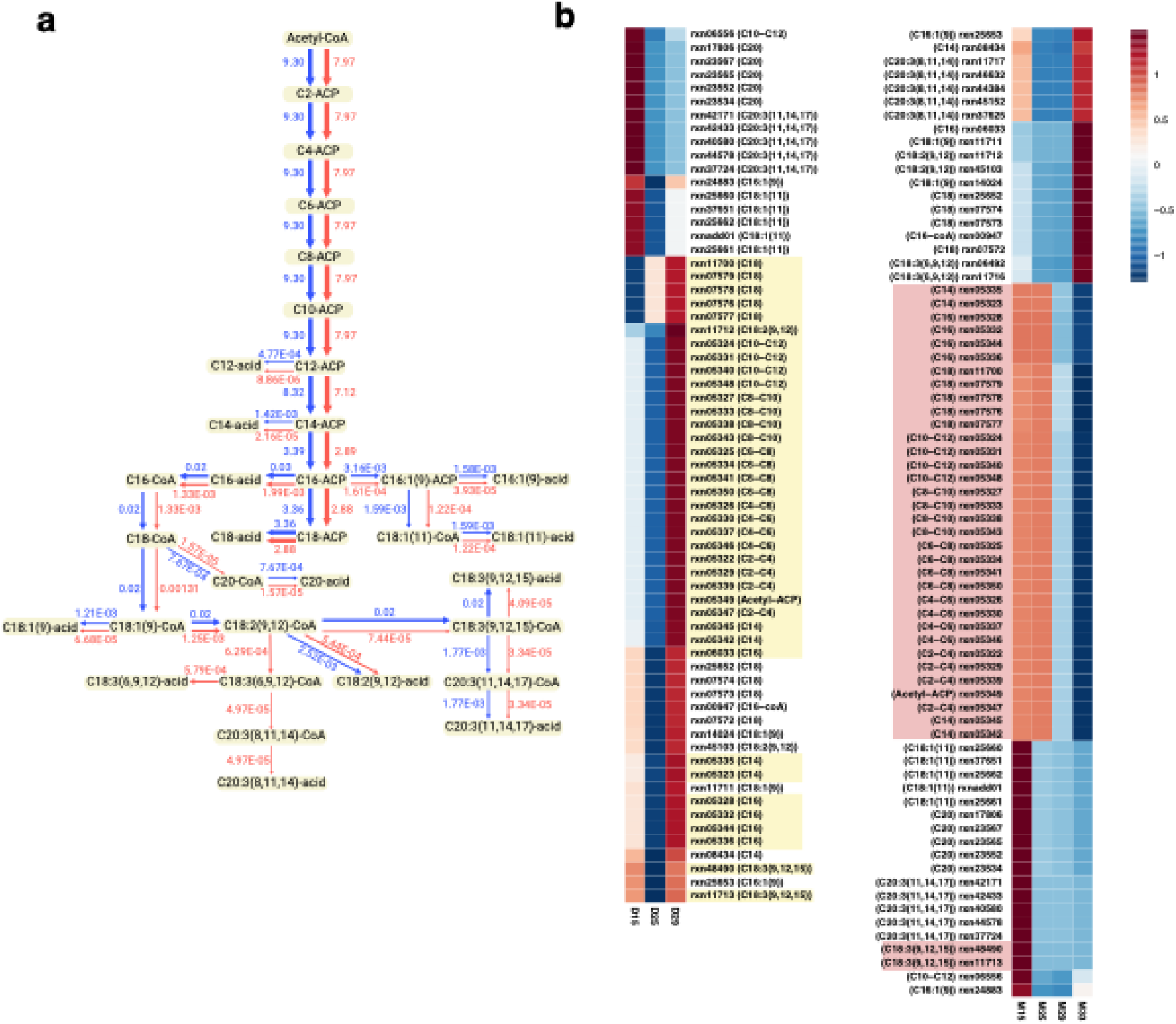
Temperature-dependent metabolic flux in fatty acid biosynthesis. **(a)** Genome-scale metabolic model-predicted fluxes through fatty acid biosynthetic reactions in *Dolichospermum* and *Microcystis aeruginosa* at their respective optimal temperatures (red: *Microcystis aeruginosa*; blue: *Dolichospermum* sp.). Heatmap of fluxes through fatty acid biosynthetic reactions across temperatures in *Dolichospermum* (left) and *Microcystis aeruginosa* (right). Yellow and red boxes highlight reactions shared between species that exhibit opposite temperature-dependent trends.

Across all select temperatures, yellow- and red-highlighted reactions in Fig. 5b mark pathways with opposite trends between *Dolichospermum* sp. and *M. aeruginosa*. These include acetyl-ACP formation, elongation from C2 to C18 saturated fatty acids, and production of PUFAs C18:2 (9,12) and C18:3 (9,12,15). *Dolichospermum* sp. displayed higher fluxes at high temperature (29°C), whereas *M. aeruginosa* exhibited higher fluxes at low temperatures (15 and 25°C). For the other reactions in the fatty-acid biosynthesis pathway, fluxes were similar between the two species. Notably, these modeled flux patterns mirrored temperature-dependent fatty acid contents observed in Fig. 2b, supporting a mechanistic basis for the lipidomic shifts.

## Methods

### Algal Strains and Culture Conditions

This study used two Dolichospermum strains (*Dolichospermum flos-aquae* FACHB 245 and *Dolichospermum* sp. YZ) and three *Microcystis* strains (*M. aeruginosa* FACHB 1322, *M. aeruginosa* FACHB 1306, and *Microcystis flos-aquae* FACHB 927). FACHB 245, FACHB 1322, FACHB 1306, and FACHB 927 were obtained from the Freshwater Algae Culture Collection at the Institute of Hydrobiology (FACHB, Chinese Academy of Sciences), whereas *Dolichospermum* sp. YZ was isolated from Lake Tai. All strains were maintained in sterile BG-11 medium at 25°C under a 14 h light / 10 h dark photoperiod with a light intensity of 40 μmol photons m⁻² s⁻¹ and regularly transferred to fresh BG-11 medium to maintain exponential growth.

To compare the physiological responses of co-occurring bloom-forming cyanobacteria under different temperatures, we selected two strains isolated from the same freshwater system—*Dolichospermum* sp. YZ and *Microcystis aeruginosa* Taihu98, both obtained from Lake Taihu—for all subsequent experiments.

### Growth Measurements

Algal growth was monitored daily by collecting 2 mL of culture under sterile conditions and fixing samples with 2% Lugol’s solution. *Dolichospermum* filaments were dispersed into single cells using an ultrasonic disruptor, without damaging cell structures, prior to counting. Cell density was determined using a hemocytometer under a microscope. Specific growth rates (μ) were calculated as the slope of ln(cell density) versus time during the exponential growth phase. Maximum growth rates were used to define species-specific optimal temperatures for subsequent experiments.

### RNA Extraction and Transcriptome Sequencing

Algal cells of equivalent biomass were collected at each temperature. Samples were flash-frozen in liquid nitrogen and ground for total RNA extraction. RNA quality was assessed via agarose gel electrophoresis, NanoPhotometer spectrophotometry, Qubit 2.0 Fluorometry, and Agilent 2100 Bioanalyzer. mRNA was isolated by removing rRNA and fragmented into 250–300 bp fragments using NEB Fragmentation Buffer. First-strand cDNA synthesis was performed with random hexamer primers, followed by second-strand synthesis and library construction according to NEB standard or strand-specific protocols. Libraries were quantified with Qubit 2.0, assessed for insert size using Agilent 2100, and validated by qRT-PCR (effective concentration > 2 nM). Qualified libraries were pooled based on target sequencing depth and sequenced on an Illumina platform using sequencing by synthesis.

Raw sequencing data were filtered to remove adapter sequences, reads with >50% low-quality bases (Q ≤ 26), or reads containing ambiguous bases (N). Clean reads were assembled de novo using Trinity, and redundancy was removed with CD-HIT-EST (sequence identity threshold 0.95) to obtain non-redundant Unigene sequences. The longest sequences per cluster were used for downstream analyses.

Differential expression across temperatures was assessed using DESeq2 (adjusted p-value < 0.05), and GO enrichment analysis was performed with topGO using all expressed genes as the background and an enrichment significance threshold of 0.05.

### Quantification of Soluble Sugars, Total Protein, and ATP

Soluble sugars were extracted and quantified using the Plant Soluble Sugar Content Assay Kit (Boxbio, AKPL008M), following the manufacturer’s instructions. Total protein content was measured using the Pierce BCA Protein Assay Kit (Thermo Scientific) via colorimetric assay. ATP was quantified in exponential-phase cultures using the ATP Assay Kit (Beyotime, S0026) according to kit instructions, using fluorescence or colorimetric detection.

### Targeted Fatty Acid Analysis

Cells of equal biomass were collected at each temperature. Algal samples were homogenized with steel beads and extracted with 1 mL chloroform:methanol (1:1, v/v). Samples were ultrasonicated for 15 min at low temperature, incubated at –20°C for 15 min, and centrifuged at 13,000 × g at 4°C for 10 min. The supernatant was dried under nitrogen, derivatized with methylation reagents (0.2 mL), incubated at 60°C for 30 min, and extracted with hexane for GC-MS analysis.

Fatty acids were separated on an Agilent DB-FastFAME capillary column (20 m × 0.18 mm × 0.20 μm) with helium as the carrier gas (1 mL min⁻¹). The column temperature program started at 80°C (30 s), ramped to 180°C at 70°C/min, then to 220°C at 4°C/min, and held at 240°C for 2 min. Samples (1 μL) were injected in split mode (50:1). Detection was by EI at 70 eV using Agilent 8890B GC coupled with 5977B MSD. Absolute concentrations were calculated using external calibration curves from peak areas.

### 13C Metabolic Flux Analysis

Exponentially growing cultures were supplemented with ¹³C-labeled sodium bicarbonate, incubated for 1 h, and sampled at equivalent cell numbers. Samples were extracted with water:methanol:chloroform (1:1:2, v/v/v), ultrasonicated on ice for 5 min, centrifuged at 3,500 × g, and the chloroform phase dried under nitrogen. Samples were hydrolyzed with 75% ethanol containing 0.5 M KOH at 80°C for 1 h, extracted with hexane, dried, and derivatized using HoBt, cholamine, and HATU. A pooled QC sample was prepared by combining equal volumes of all samples.

Chromatography was performed on a ThermoFisher Ultimate 3000 UHPLC system with Waters BEH C18 column (2.1 × 100 mm, 1.7 μm) at a flow rate of 0.25 mL/min. Mobile phases were water (A) and acetonitrile (B) with 0.1% formic acid. Linear gradient elution was applied (0–18 min, 90–0% B). Mass spectrometry was performed on a Thermo Q Exactive Orbitrap in negative HESI mode. Raw data were integrated and corrected for natural isotope abundance using published methods.

### Fatty Acid Supplementation Experiments

Fatty acids showing the largest changes at non-optimal temperatures relative to the optimum were selected for supplementation. The amount added was calculated based on measured intracellular concentrations and daily cell counts. Exogenous fatty acids were added daily for 5 days at concentrations corresponding to physiological levels, with 10× and 100× dilutions to avoid excessive supplementation. Control cultures received no fatty acids. Cell densities were measured daily, and only the concentration showing the strongest effect was reported.

### Molecular Evolution Analyses

Core metabolic and fatty acid biosynthetic genes from 52 *Dolichospermum* and 50 Microcystis genomes were analyzed. Protein sequences of orthologs were aligned, and the ratio of nonsynonymous to synonymous substitutions(dN/dS) was calculated using PAML to assess evolutionary conservation.

### Temperature-Responsive Regulatory Mechanisms

Known transcription factors NtcA and LexA and their downstream targets (*gln*B, *nir*A, *gln*A, *hox*E) were identified in both species via BLAST. Expression patterns were visualized as heatmaps to examine temperature-dependent regulation.

The thermosensor Hik34 and its putative downstream regulator Rre1 were identified through BLAST. Promoter regions of *rpo*D and candidate downstream genes were extracted and scanned with a sliding window (20 bp, step 5 bp) to avoid length biases. Protein–DNA interactions between Rre1 or RpoD and promoter fragments were predicted using AlphaFold3^18^ with predicted TM-score (ptm) > 0.5 and interface pTM (ipTM) > 0.6 considered high-confidence.

### Genome-Scale Metabolic Modeling and Flux Simulations

Draft genome-scale metabolic models for *Dolichospermum* sp. and *Microcystis aeruginosa* were generated using ModelSEED^20^ and converted to enzyme-constrained models with GECKO^19^.

Temperature-specific gene expression ratios (low or high relative to the optimum) were applied to adjust enzyme constraints for each reaction. For reactions controlled by multiple genes, enzyme constraints were adjusted based on gene logic: the maximal fold-change was applied for OR-linked genes, whereas the minimal fold-change was used for AND-linked genes.Experimentally measured fatty acid compositions were incorporated into biomass reactions as absolute molar concentrations (mmol/gDW). Flux balance analysis was performed for each temperature condition with biomass production as the objective, and fluxes through fatty acid biosynthesis were extracted to quantify temperature-dependent shifts in lipid metabolism.

## Discussion

Climate change and recurrent heatwaves have elevated temperature to a ‘co-equal’ determinant of CHABs alongside continued nutrient loading. Yet, despite its central importance, the thermal dimension of CHAB ecology remains poorly understood. Here we define the complete thermal niches of two globally dominant bloom-forming cyanobacteria and elucidate the systems-level mechanisms that underpin their separate temperature optima, which is instrumental in both bloom control and general climate change studies.

The thermal separation between filamentous (often nitrogen-fixing) cyanobacteria, e.g., *Aphanizomenon* spp. and *Dolichospermum* spp. and colonial *M. aeruginosa* has been fairly studied different in many ways ^12,21–23^. The former grows better at low temperatures and the latter grows better at high temperatures. Our study not only confirmed this with more species but also found that these two species have overlapping thermal range, in which they grow similarly between 18 and 25°C. More importantly, we discovered that this niche separation is due to their wholistic cellular responses to temperatures. Intracellular contents of macromolecules—ATP, proteins, lipids, and carbohydrates— are different between the two species. Additionally, gene expression, fatty acid composition, and metabolic fluxes toward FAs are also different. Finally, we identified the FAs which are experimentally confirmed to support niche separation.

From a systems-biology perspective, this niche separation is a systems-level response to temperature. That is, cells respond by integrating external stimuli into transcriptional and metabolic regulatory signals into genome-scale metabolic network and operate toward an optimum equilibrium ^24^. One major finding is the thermal sensor Hik34, based on a homolog in *Synechococcus elongatus* PCC 7942^16^ and its entire regulatory cascade to effector genes. The homologous sensor kinase indeed phosphorylates effector protein Rre1, which binds the *rpoD* promoter regions in *Dolichospermum* sp. and *M. aeruginosa*, while expressed *σ*^70^ in turn directly controls the expression of downstream stress-responsive target genes.

Finally, we model the fluxes toward fatty acids using GSMs for each species. By reconstructing a GSM and integrating transcriptome and enzyme parameters, we obtained fluxes toward critical fatty acids observed to support thermal tolerance in each species.

To sum up, this study not only presented a complete thermal range for either of the two top blooming cyanobacteria and identified major macromolecules sustaining their niche separation. More importantly, we also delineated their regulatory hierarchy trigger by thermal stress and the systems-biology mechanisms of response based on genome-scale metabolic network. This work will prove instrumental in bloom prediction and control and more generally in climate change studies.

