## Supplementary Figure 1 for "Thermal boundaries and underlying systems biology mechanisms in two waterbloom cyanobacteria"

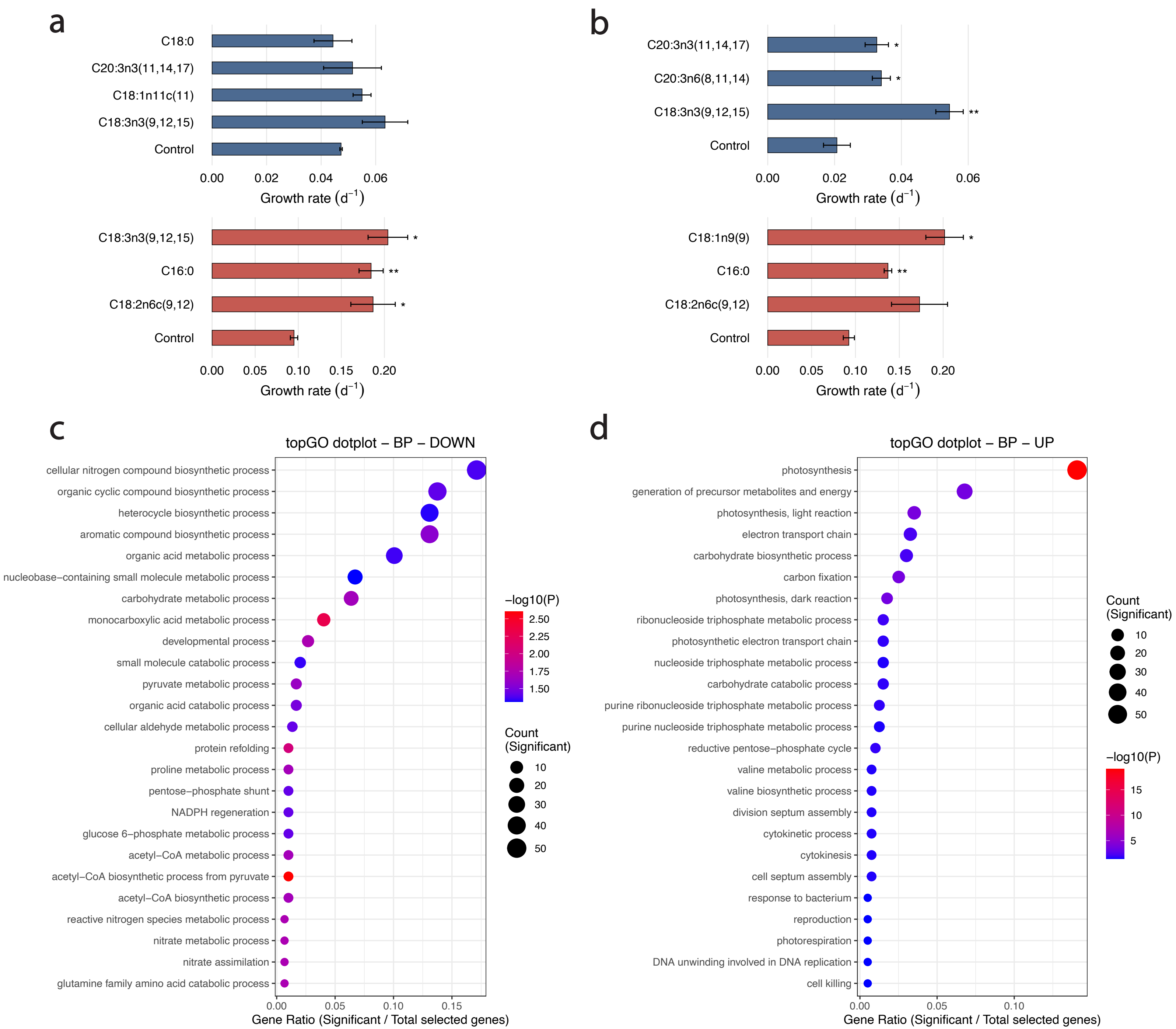
a
C18:0
C20:3n3(11,14,17)
C18:1n11c(11)
C18:3n3(9,12,15)
Control
0.00 0.02 0.04 0.06
Growth rate (d$^{-1}$)
C18:3n3(9,12,15) *
C16:0 **
C18:2n6c(9,12) *
Control
0.00 0.05 0.10 0.15 0.20
Growth rate (d$^{-1}$)
b
C20:3n3(11,14,17) *
C20:3n6(8,11,14) *
C18:3n3(9,12,15) **
Control
0.00 0.02 0.04 0.06
Growth rate (d$^{-1}$)
C18:1n9(9) *
C16:0 **
C18:2n6c(9,12)
Control
0.00 0.05 0.10 0.15 0.20
Growth rate (d$^{-1}$)
c
topGO dotplot – BP – DOWN
cellular nitrogen compound biosynthetic process
organic cyclic compound biosynthetic process
heterocycle biosynthetic process
aromatic compound biosynthetic process
organic acid metabolic process
nucleobase−containing small molecule metabolic process
carbohydrate metabolic process
monocarboxylic acid metabolic process
developmental process
small molecule catabolic process
pyruvate metabolic process
organic acid catabolic process
cellular aldehyde metabolic process
protein refolding
proline metabolic process
pentose−phosphate shunt
NADPH regeneration
glucose 6−phosphate metabolic process
acetyl−CoA metabolic process
acetyl−CoA biosynthetic process from pyruvate
acetyl−CoA biosynthetic process
reactive nitrogen species metabolic process
nitrate metabolic process
nitrate assimilation
glutamine family amino acid catabolic process
0.00 0.05 0.10 0.15
Gene Ratio (Significant / Total selected genes)
−log10(P)
2.50
2.25
2.00
1.75
1.50
Count (Significant)
10
20
30
40
50
d
topGO dotplot – BP – UP
photosynthesis
generation of precursor metabolites and energy
photosynthesis, light reaction
electron transport chain
carbohydrate biosynthetic process
carbon fixation
photosynthesis, dark reaction
ribonucleoside triphosphate metabolic process
photosynthetic electron transport chain
nucleoside triphosphate metabolic process
carbohydrate catabolic process
purine ribonucleoside triphosphate metabolic process
purine nucleoside triphosphate metabolic process
reductive pentose−phosphate cycle
valine metabolic process
valine biosynthetic process
division septum assembly
cytokinetic process
cytokinesis
cell septum assembly
response to bacterium
reproduction
photorespiration
DNA unwinding involved in DNA replication
cell killing
0.00 0.05 0.10
Gene Ratio (Significant / Total selected genes)
Count (Significant)
10
20
30
40
50
−log10(P)
15
10
5

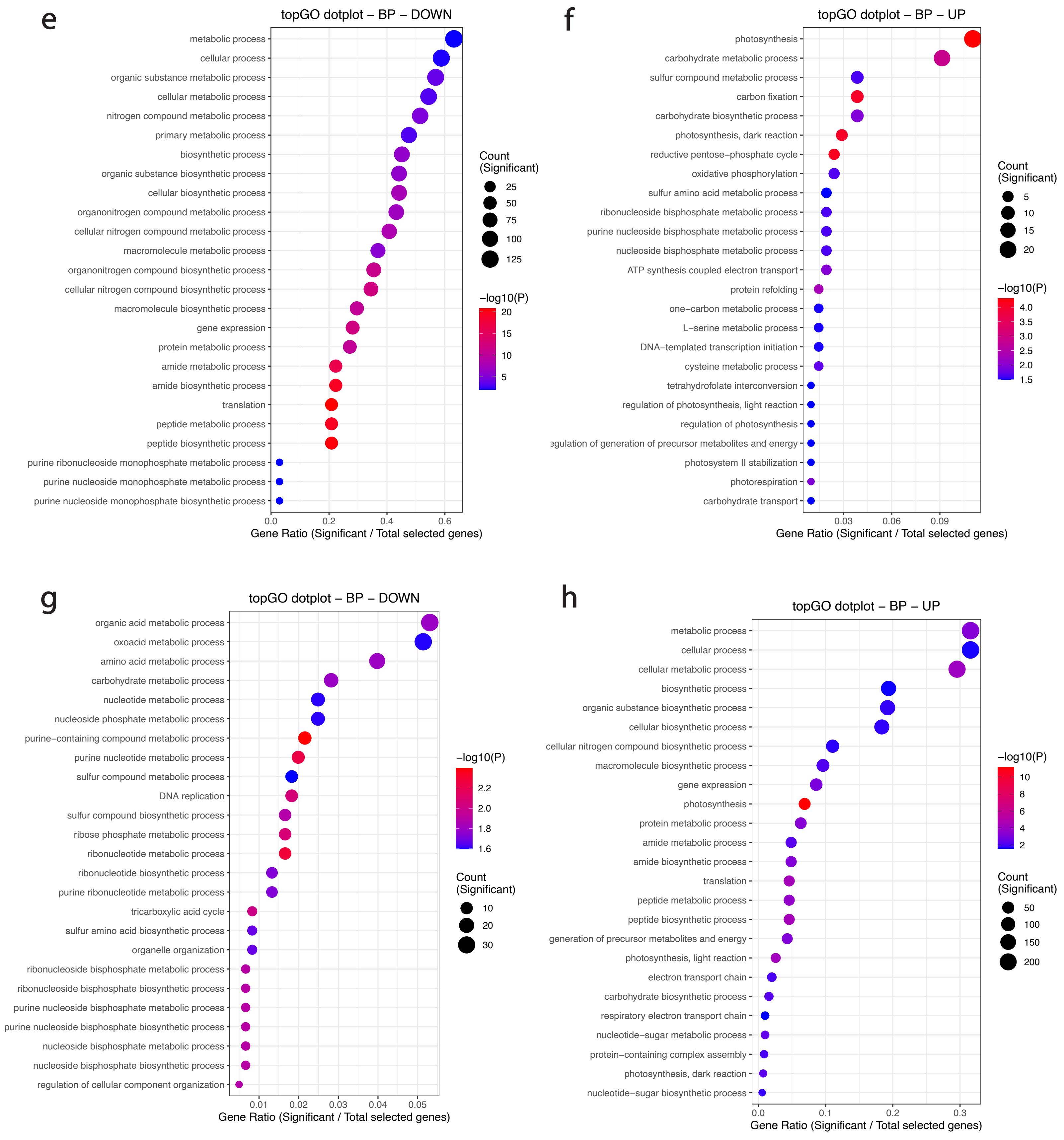
e
topGO dotplot – BP – DOWN
metabolic process
cellular process
organic substance metabolic process
cellular metabolic process
nitrogen compound metabolic process
primary metabolic process
biosynthetic process
organic substance biosynthetic process
cellular biosynthetic process
organonitrogen compound metabolic process
cellular nitrogen compound metabolic process
macromolecule metabolic process
organonitrogen compound biosynthetic process
cellular nitrogen compound biosynthetic process
macromolecule biosynthetic process
gene expression
protein metabolic process
amide metabolic process
amide biosynthetic process
translation
peptide metabolic process
peptide biosynthetic process
purine ribonucleoside monophosphate metabolic process
purine nucleoside monophosphate metabolic process
purine nucleoside monophosphate biosynthetic process
0.0
0.2
0.4
0.6
Gene Ratio (Significant / Total selected genes)
Count (Significant)
25
50
75
100
125
−log10(P)
20
15
10
5
f
topGO dotplot – BP – UP
photosynthesis
carbohydrate metabolic process
sulfur compound metabolic process
carbon fixation
carbohydrate biosynthetic process
photosynthesis, dark reaction
reductive pentose−phosphate cycle
oxidative phosphorylation
sulfur amino acid metabolic process
ribonucleoside bisphosphate metabolic process
purine nucleoside bisphosphate metabolic process
nucleoside bisphosphate metabolic process
ATP synthesis coupled electron transport
protein refolding
one−carbon metabolic process
L−serine metabolic process
DNA−templated transcription initiation
cysteine metabolic process
tetrahydrofolate interconversion
regulation of photosynthesis, light reaction
regulation of photosynthesis
egulation of generation of precursor metabolites and energy
photosystem II stabilization
photorespiration
carbohydrate transport
0.03
0.06
0.09
Gene Ratio (Significant / Total selected genes)
Count (Significant)
5
10
15
20
−log10(P)
4.0
3.5
3.0
2.5
2.0
1.5
g
topGO dotplot – BP – DOWN
organic acid metabolic process
oxoacid metabolic process
amino acid metabolic process
carbohydrate metabolic process
nucleotide metabolic process
nucleoside phosphate metabolic process
purine−containing compound metabolic process
purine nucleotide metabolic process
sulfur compound metabolic process
DNA replication
sulfur compound biosynthetic process
ribose phosphate metabolic process
ribonucleotide metabolic process
ribonucleotide biosynthetic process
purine ribonucleotide metabolic process
tricarboxylic acid cycle
sulfur amino acid biosynthetic process
organelle organization
ribonucleoside bisphosphate metabolic process
ribonucleoside bisphosphate biosynthetic process
purine nucleoside bisphosphate metabolic process
purine nucleoside bisphosphate biosynthetic process
nucleoside bisphosphate metabolic process
nucleoside bisphosphate biosynthetic process
regulation of cellular component organization
0.01
0.02
0.03
0.04
0.05
Gene Ratio (Significant / Total selected genes)
−log10(P)
2.2
2.0
1.8
1.6
Count (Significant)
10
20
30
h
topGO dotplot – BP – UP
metabolic process
cellular process
cellular metabolic process
biosynthetic process
organic substance biosynthetic process
cellular biosynthetic process
cellular nitrogen compound biosynthetic process
macromolecule biosynthetic process
gene expression
photosynthesis
protein metabolic process
amide metabolic process
amide biosynthetic process
translation
peptide metabolic process
peptide biosynthetic process
generation of precursor metabolites and energy
photosynthesis, light reaction
electron transport chain
carbohydrate biosynthetic process
respiratory electron transport chain
nucleotide−sugar metabolic process
protein−containing complex assembly
photosynthesis, dark reaction
nucleotide−sugar biosynthetic process
0.0
0.1
0.2
0.3
Gene Ratio (Significant / Total selected genes)
−log10(P)
10
8
6
4
2
Count (Significant)
50
100
150
200

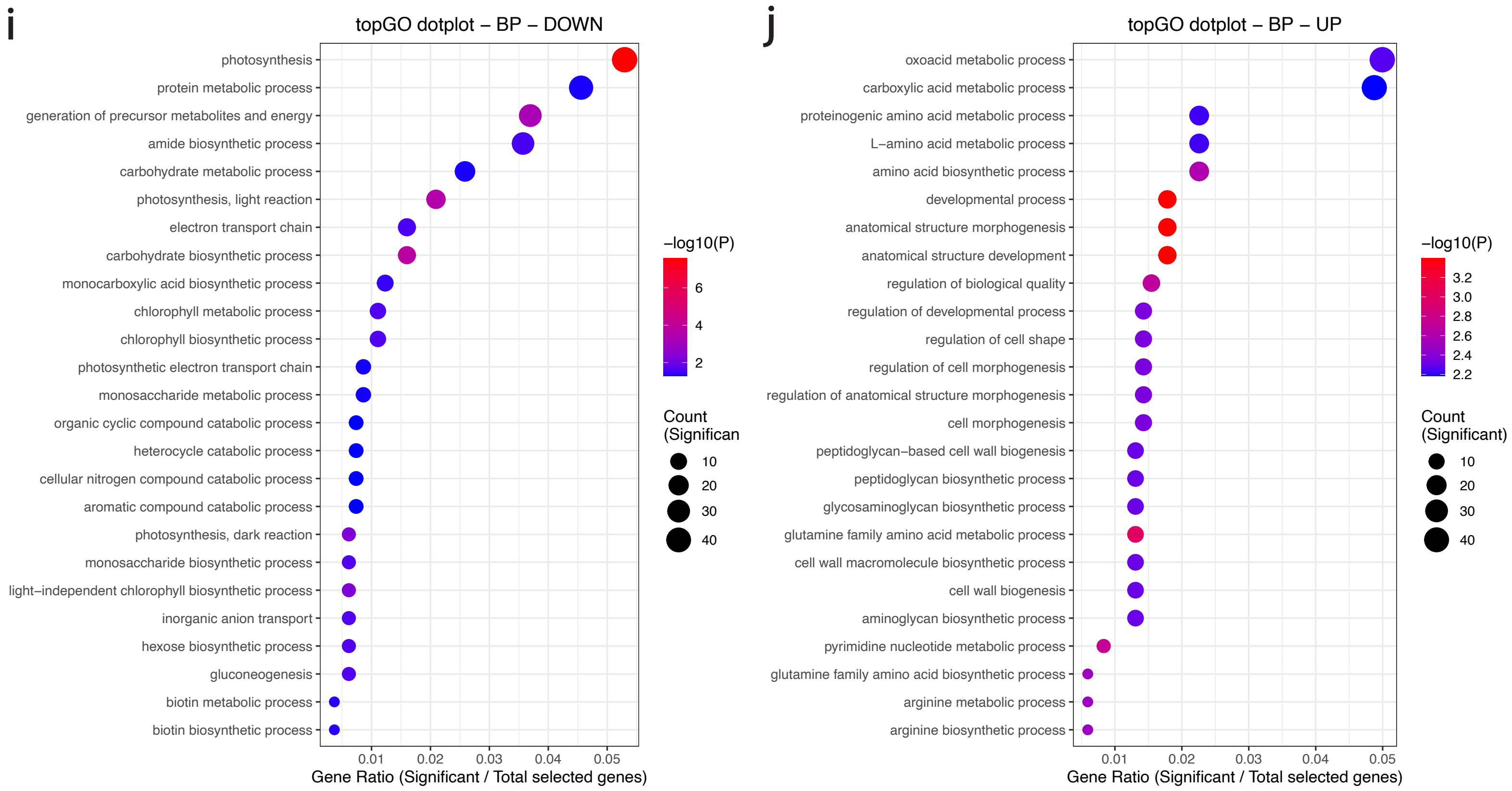


**Supplementary Figure 1. Growth responses and functional enrichment.**

(a) Maximum growth rates of *Dolichospermum* with exogenous fatty acid supplementation under low (blue) and high (red) temperatures.
(b) Maximum growth rates of *Microcystis aeruginosa* with exogenous fatty acid supplementation under low (blue) and high (red) temperatures.
(c,d) Functional enrichment of downregulated (left) and upregulated (right) genes in *Dolichospermum* at 15 °C relative to 25 °C.
(e,f) Functional enrichment of downregulated (left) and upregulated (right) genes in *Dolichospermum* at 29 °C relative to 25 °C.
(g,h) Functional enrichment of downregulated (left) and upregulated (right) genes in *Microcystis aeruginosa* at 15 °C relative to 29 °C.
(i,j) Functional enrichment of downregulated (left) and upregulated (right) genes in *Microcystis aeruginosa* at 33 °C relative to 29 °C.
